# *Notch1* is required to maintain supporting cell identity and vestibular function during maturation of the mammalian balance organs

**DOI:** 10.1101/2024.06.21.600098

**Authors:** Alison Heffer, Choongheon Lee, Joseph C. Holt, Amy E. Kiernan

## Abstract

The inner ear houses two sensory modalities: the hearing organ, located in the cochlea, and the balance organs, located throughout the vestibular regions of the ear. Both hearing and vestibular sensory regions are composed of similar cell types, including hair cells and associated supporting cells. Recently, we showed that *Notch1* is required for maintaining supporting cell survival postnatally during cochlear maturation. However, it is not known whether *Notch1* plays a similar role in the balance organs of the inner ear. To characterize the role of Notch during vestibular maturation, we conditionally deleted *Notch1* from *Sox2*-expressing cells of the vestibular organs in the mouse at P0/P1. Histological analyses showed a dramatic loss of supporting cells accompanied by an increase in type II hair cells without cell death, indicating the supporting cells are converting to hair cells in the maturing vestibular regions. Analysis of 6-week old animals indicate that the converted hair cells survive, despite the reduction of supporting cells. Interestingly, measurements of vestibular sensory evoked potentials (VsEPs), known to be generated in the striolar regions of the vestibular afferents in the maculae, failed to show a response, indicating that NOTCH1 expression is critical for striolar function postnatally. Consistent with this, we find that the specialized type I hair cells in the striola fail to develop the complex calyces typical of these cells. These defects are likely due to the reduction in supporting cells, which have previously been shown to express factors critical for the striolar region. Similar to other mutants that lack proper striolar development, *Notch1* mutants do not exhibit typical vestibular behaviors such as circling and head shaking, but do show difficulties in some vestibular tests, including the balance beam and forced swim test. These results indicate that, unlike the hearing organ in which the supporting cells undergo cell death, supporting cells in the balance regions retain the ability to convert to hair cells during maturation, which survive into adulthood despite the reduction in supporting cells.

**Significance Statement:** Notch signaling regulates the cell fate choices between hair cells and supporting cells during inner ear development. However, little is known about how Notch functions in the mammalian vestibular sensory organs once cell fate has been determined. Here, we examine the role of *Notch1* in the maturing balancing organs. We show that deletion of *Notch1* results in vestibular physiological and behavioral dysfunction by 3 months of age. Histological analyses reveal supporting cells are converting to type II hair cells in the utricle, and despite a loss of supporting cells, the hair cells survive to adulthood. Additionally, the striolar type I hair cells important for generating a VsEP response are decreased in number and not innervated properly. These results show that Notch continues to function in maintaining supporting cell identity in the vestibular organs postnatally, which may be important in strategies for hair cell regeneration.

## Introduction

The mammalian inner ear is comprised of the cochlea and vestibular end organs, a collection of sensory organs that mediate hearing and balance, respectively. The sensory neuroepithelium of the vestibular end organs is represented by two otolithic maculae – the saccule and utricle – which detect linear acceleration, and three canal cristae that detect angular acceleration (Burns and Stone, 2017; Mackowetzky et al., 2021). Deficits or dysfunction in one or more regions of the vestibular system can lead to balance issues, dizziness, and vertigo (Mackowetzky et al., 2021). However, the underlying cellular deficits or problems associated with many of these vestibular behaviors are still not well understood.

Within each vestibular epithelium, there are sensory hair cells that detect head movements, as well as non-sensory supporting cells that surround the hair cells. The hair cells are innervated by both afferent and efferent vestibular neurons that transmit information to and from the brain (Wan et al., 2013; Burns and Stone, 2017). All balance organs contain two main types of hair cells which can be classified by their characteristic morphology, gene expression profile, type of afferent innervation and physiological responses (Eatock et al., 1998; Rusch et al., 1998; Moravec and Peterson, 2004; Li et al., 2008; Eatock and Songer, 2011; Deans, 2013; Burns and Stone, 2017; McInturff et al., 2018). Type I hair cells are flask-shaped with long, thick stereocilia when mature, and surrounded by afferent calyces. Type II hair cells are cylindrical in shape with shorter, thinner stereocilia, and are innervated as afferent boutons. Additionally, the striola of the maculae balance organs and central zone of the cristae are enriched with a subset of specialized type I hair cells. In maculae, these cells are innervated by complex calyces, and are known to be important in generating the vestibular sensory evoked potential (VsEP) response (Ono et al., 2020; Kim et al., 2022).

Supporting cells of the inner ear have many important functions, including structural support, maintaining ion and small molecule homeostasis, clearing cellular debris, and maintaining hair cell survival (Monzack and Cunningham, 2013; Wan et al., 2013). While supporting cell subtypes, their overall appearance, and their gene expression profiles are quite diverse in the cochlea, supporting cells within the vestibular sensory regions are generally more homogeneous in appearance, though little is known about the diversity across the sensory epithelium. However, differences do exist, as studies looking at different Cre lines and their efficiency in the adult utricle found that while some Cre lines target all supporting cells in the utricle, other Cre lines target either extrastriolar or striolar supporting cells more efficiently (Burns et al., 2012; Stone et al., 2018; You et al., 2018). Single-cell RNA-seq data from both the entire utricular sensory epithelium and from extrastriolar and striolar supporting cells have also revealed distinct populations and gene expression profiles (You et al., 2018; Jan et al., 2021).

Notch signaling plays a pivotal role during inner ear development by regulating several developmental processes including cell proliferation, cell fate acquisition and cell survival (Radtke et al., 2010; Kiernan, 2013; Heffer et al., 2023). During inner ear embryogenesis, Notch functions through canonical signaling, via the binding of a Notch ligand on one cell (JAGGED 1, 2 or DLL 1,3, 4) with a Notch receptor (NOTCH1-4) on a neighboring cell, resulting in the cleavage of the Notch intracellular domain (NICD) and its translocation to the nucleus where Notch targets are activated or repressed (Lanford et al., 1999; Mumm and Kopan, 2000; Kopan and Ilagan, 2009). Notch is initially required for development of the sensory regions via its ligand Jagged1 (JAG1) (Kiernan et al., 2001; Kiernan et al., 2006; Saravanamuthu et al., 2009; Neves et al., 2011; Brown et al., 2020), through a process known as lateral induction. Subsequently, Notch signaling is also essential for the determination of hair cell or supporting cell fate via a process known as lateral inhibition. Here, activation of Notch via the ligand on a neighboring cell leads to inhibition of the hair cell fate, and acquisition of the supporting cell fate. Thus, a mosaic of cell types is produced in which each hair cell is surrounded by supporting cells. Evidence for this mechanism comes from experiments in which dysfunction of Notch signaling leads to an increase in hair cells at the expense of supporting cells (Lanford et al., 1999; Zine et al., 2000; Kiernan et al., 2005; Kiernan et al., 2006). Once hair cells and supporting cells have adopted their fate and begin differentiating, the role of Notch is less clear, particularly the role of specific Notch ligands and receptors.

Previously, we have shown in the postnatal cochlea, that deletion of the *Notch1* receptor results in rapid death of the supporting cells, leading secondarily to loss of hair cells and deafness (Heffer et al., 2023). Similarly, loss of the Notch ligand JAG1 in the postnatal cochlea leads to loss of the Hensen’s supporting cells and defects in stereocilia maturation (Chrysostomou et al., 2020; Gilels et al., 2022). Additionally, *in vitro* experiments in utricle cultures, in which Notch signaling was inhibited though DAPT treatment, suggested a role for Notch in supporting cell proliferation and differentiation (Wu et al., 2016). However, it is not known what role Notch signaling plays during maturation of the vestibular sensory regions *in vivo*, particularly what role specific Notch ligands and receptors play. Moreover, fewer studies have correlated the effects of cellular changes on vestibular function.

Here, we conditionally delete *Notch1* at birth (P0/P1) in all *Sox2*-expressing cells of the vestibular regions and examine both the functional and cellular consequences of loss of Notch signaling during vestibular maturation. We find that while there are no obvious behaviors previously associated with vestibular defects like circling or head bobbing, *Notch1*-deficient mice exhibited balance and swimming problems, suggesting that *Notch1* deletion results in vestibular impairment. In agreement with this, a test of macular function using linear head motions, the VsEP, shows significantly compromised vestibular function in *Notch1*-deleted animals. Further immunohistochemical analyses revealed a significant loss of supporting cells of all vestibular sensory regions at 6 weeks, and a dramatic increase in type II hair cells. Lack of Caspase-3+ staining indicates that, unlike the cochlea, these supporting cells are not dying but are instead transdifferentiating directly into hair cells. Consistent with this, an up-regulation of the transcription factor ATOH1, a gene required for hair cell development, is observed in many supporting cells in the *Notch1*-deficient utricle. Additionally, *Notch1* mutants have abnormalities in the striolar region: there is a significant loss of specialized type I hair cells from the striolar region and abnormal innervation of those remaining. These studies reveal that the maturing vestibular system responds differently to loss of NOTCH1 than the maturing cochlea. However, reduction of the supporting cell population, and/or increase in hair cells leads to vestibular dysfunction despite the lack of cell death, suggesting NOTCH1 is important for maintaining integrity of the balance organs postnatally.

## Materials and Methods

### Animals and tamoxifen treatment

All procedures in this study were performed in accordance with guidelines and regulations of the University of Rochester Medical Center and the Guide for the Care and Use of Laboratory Animals of the National Institutes of Health, and all animal experiments were approved by the University of Rochester Committee on Animal Resources. The mouse strains used in these studies were: *Sox2-CreER* (Stock #017593, The Jackson Laboratory; (Arnold et al., 2011)) and *Notch1^flox^* (a gift from Raphael Kopan; (Yang et al., 2004)). Animals were genotyped using the following primers: Sox2Cre1: 5’-TGA TGA GGT TCG CAA GAA CC and Sox2Cre2 5’-CCA TGA GTG AAC GAA CCT GG (yielding a 350 bp product) and Notch1F: 5’-TGC CCT TTC CTT AAA AGT GG and Notch1R: 5’-GCC TAC TCC GAC ACC CAA TA (mutants had a 281 bp product). Tamoxifen (Sigma-Aldrich) was dissolved in corn oil and injected within 4 hours of birth (P0) and 20-24 hours later (P1); pups were given intramuscular injections of tamoxifen (37.5 µg/g body weight) into the muscle on the left leg, parallel to vein. Animals that were used for uninjected controls came from litters that we born and aged the same as those that received tamoxifen injection. Both male and female mice were used in this study.

### Vestibular Stimulus Evoked Potentials (VsEPs)

All physiological recordings were conducted in a sound-treated, electrically shielded booth. Anesthesia was induced in all mice using intraperitoneal urethane/xylazine (1.2 g/kg/20 mg/kg). Under anesthesia, body temperature was maintained at 38.0 ± 0.2°C, monitored via noninvasive rectal temperature measurement. Subcutaneous electrodes were placed on the midline over the nuchal crest (G1), behind the right pinna (G2), and on the belly (GND). To ensure the absence of auditory response during VsEP testing, a binaural forward masker (intensity: 92 dB SPL; bandwidth: 50∼50 kHz) was presented using a free-field speaker (Tucker-Davis Technologies, MF1, Alachua, FL). VsEPs were amplified by a factor of 200,000 and filtered between 0.3-3 kHz at −6 dB amplitude points (Grass P511, Model K). Electrophysiological signals were analog-to-digital converted, with 1,024 points sampled at 10 µs/point, triggered at the onset of each stimulus. Signal averaging was employed to extract responses to from background noise, generating VsEP response waveforms. Each VsEP waveform was constructed from separate average responses to each polarity (256 sweeps each), totaling 512 sweeps.

To secure the mouse’s head to an electromechanical shaker, a noninvasive head clip was utilized (Jones et al., 2015). The electromechanical shaker (Labworks Inc., ET-132-2, Costa mesa, CA) delivers a linear voltage ramp along the naso-occipital axis (± X), directly stimulating otolith organs. An accelerometer (100 mV/g, g = 9.81 m/s^2^) mounted on the shaker platform monitored jerk magnitude. Stimulus level, measured as mean peak jerk amplitude, was expressed in dB relative to 1 g/ms. A threshold-seeking protocol was implemented, adjusting stimulus level in 3-dB steps within a range of +6 dB to −18 dB re: 1.0 g/ms. VsEP response parameters analyzed included latencies (P1 and N1), amplitudes (P1-N1), and threshold. P1 and N1 peaks reflect the compound action potential of the peripheral vestibular nerve innervating macular sensors (Jones et al., 1999; Jones and Jones, 1999). VsEP threshold indicates the sensitivity of peripheral macular sensors to transient linear head acceleration.

### Vestibular behavior testing

#### Open field

Animals were placed in the center of a ∼50cm circular ring and movement was recorded for 15 minutes using Ethovision15 software. Movement plots as well as total distance and velocity were determined by EthoVision15 software and plotted using Prism.

#### Balance beam

Animals were tested for three consecutive days: the first two on a 20mm balance beam and the third on a 9mm bar. A dark enclosure was placed at the end of the bar for animals to use as a safe space, and a hammock sheet was placed below the balance beam to catch the animals if they fell. Animals were placed on one end of the bar and the time it took the animal to cross 80 cm to the other side was recorded on the third day (9mm bar). If animals failed to traverse the 80 cm distance in two minutes, they were given a time of 120 seconds. Behaviors associated with balance problems (legs slipping on bar, tail wrapping around bar, head tilt) were also recorded if present.

#### Swim

3.5 liters of water (77-81°F) was placed in a clear circular container 10 inches in diameter. Animals were carefully placed into the water to ensure their heads were above at the start of the swim test and carefully monitored for 4 minutes. If animals were able to keep their head and nose above water for the duration of the swim test, they were given a swim test time of 240 seconds. For animals that failed to keep their head and nose above water, the time given for the swim test was marked as when the nose first went underwater.

### Utricle fixation and Immunohistochemistry

#### Fixation

The ear was dissected out of the head at time-points of interest and fixed in 4% paraformaldehyde (Santa Cruz) overnight at 4°C. If the ear bone had undergone calcification (older than P6), samples were incubated in 0.2M EDTA for 1-2 weeks.

#### Immunofluorescence and confocal imaging

Utricles were dissected out of the ear, washed in PBS, and then transferred to 30% sucrose and flash frozen in liquid nitrogen for antigen retrieval. After thawing, utricles were washed in PBSTx (0.2% Triton-X), blocked for 2 hours in 10% horse serum (Sigma-Aldrich)/PBSTx and then incubated overnight in primary antibody at 4°C. After washing several times in PBSTx, samples were incubated in secondary antibodies for 2 hours at room temperature, washed, and mounted in Fluoro-Gel (Electron Microscopy Sciences). Confocal imaging was done using a Nikon A1-R confocal microscope with Nikon NIS-Elements software.

#### Paraffin sections and staining

After fixation and decalcification, ear tissue was dehydrated with a series of EtOH washes from 30% to 100%, cleared in xylenes, and embedded in paraffin. Paraffin blocks were sectioned at a thickness of 10 µm using a Microm HM310 microtome and dried on Superfrost Plus slides (Fisher Scientific). For antibody staining of the vestibular sensory regions, slides were deparaffinized and rehydrated through a series of xylene/ethanol washes, and then antigen retrieval was performed by incubating in slides in boiling 10mM sodium citrate buffer, pH 6, for 10-15 min. Tissue was blocked with 10% horse serum/PBS for 1–2 h and then incubated in primary antibodies in PBS/5% horse serum overnight at 4°C. The following day the slides were washed, tissue was incubated in secondary antibodies for 2 h at room temperature, and then slides were washed and mounted in Fluoro-Gel. All images were taken on a Zeiss Axio microscope using AxioVision SE64 software.

#### Antibodies

Primary antibodies used in this study were: anti-Notch1 (rabbit, 1:250; Abcam), anti-Sox2 (goat, 1:800; Santa Cruz Biotechnology), anti-Sox9 (rabbit, 1:500, Millipore Sigma), anti-Myo7a (rabbit, 1:1000; Proteus), anti-cleaved-Casp3 (rabbit, 1:1000; R&D Systems), anti-Atoh1 (rabbit, 1:50; ProteinTech), anti-Spp1(OPN) (mouse, 1:300, Santa Cruz Biotechnology), anti-Calb2 (rabbit, 1:500, Millipore Sigma), anti-TUJ (mouse, 1:1000, BioLegend), Annexin A4 (goat, 1:500; R&D Systems), and anti-Ocm (goat, 1:250, Invitrogen). Secondary antibodies used were: Alexa Fluor 555 donkey anti-mouse (1:500; Invitrogen), Alexa Fluor 488 donkey anti-rabbit (1:500; Invitrogen), Alexa Fluor 647 donkey anti-rabbit (1:500; Invitrogen), Alexa Fluor 546 donkey anti-goat (1:500; Invitrogen), Alexa Fluor 647 donkey anti-mouse (1:500; Invitrogen), and Alexa Fluor 488 donkey anti-goat (1:500; Invitrogen).

### Quantification and Statistical Analyses

Significant differences in VsEP thresholds and behavior tests were determined through two-way ANOVA tests, followed by Tukey’s post hoc analyses using Prism software. All p values <0.05 were considered significant. Quantification of all supporting cells and hair cells were performed by counting the number of cells in 10um x 15um (15,000um^2^) region of the striola and extrastriolar regions. Both one-way and two-way ANOVA with Tukey’s post hoc test were used to determine significant differences in cell counts. For all, a p value of <0.05 was considered significant.

## Results

### Notch1 is expressed in vestibular supporting cells and is required for vestibular function

To examine the role of *Notch1* in the postnatal vestibular system, we crossed tamoxifen-inducible *Sox2^CreERT2^* mice (Arnold et al., 2011) with floxed *Notch1* allele mice (Yang et al., 2004) to conditionally delete *Notch1* in the vestibular sensory regions at P0/P1, and harvested tissues at set time-points after (Figure 1A). Similar to NOTCH1 expression in the cochlea (Maass et al., 2015; Heffer et al., 2023), NOTCH1 was localized to the membrane of all vestibular supporting cells and their projections, and we find that two tamoxifen injections (P0 and P1) are sufficient to reduce NOTCH1 protein levels by P6 (Figure 1B).

**Figure 1.**
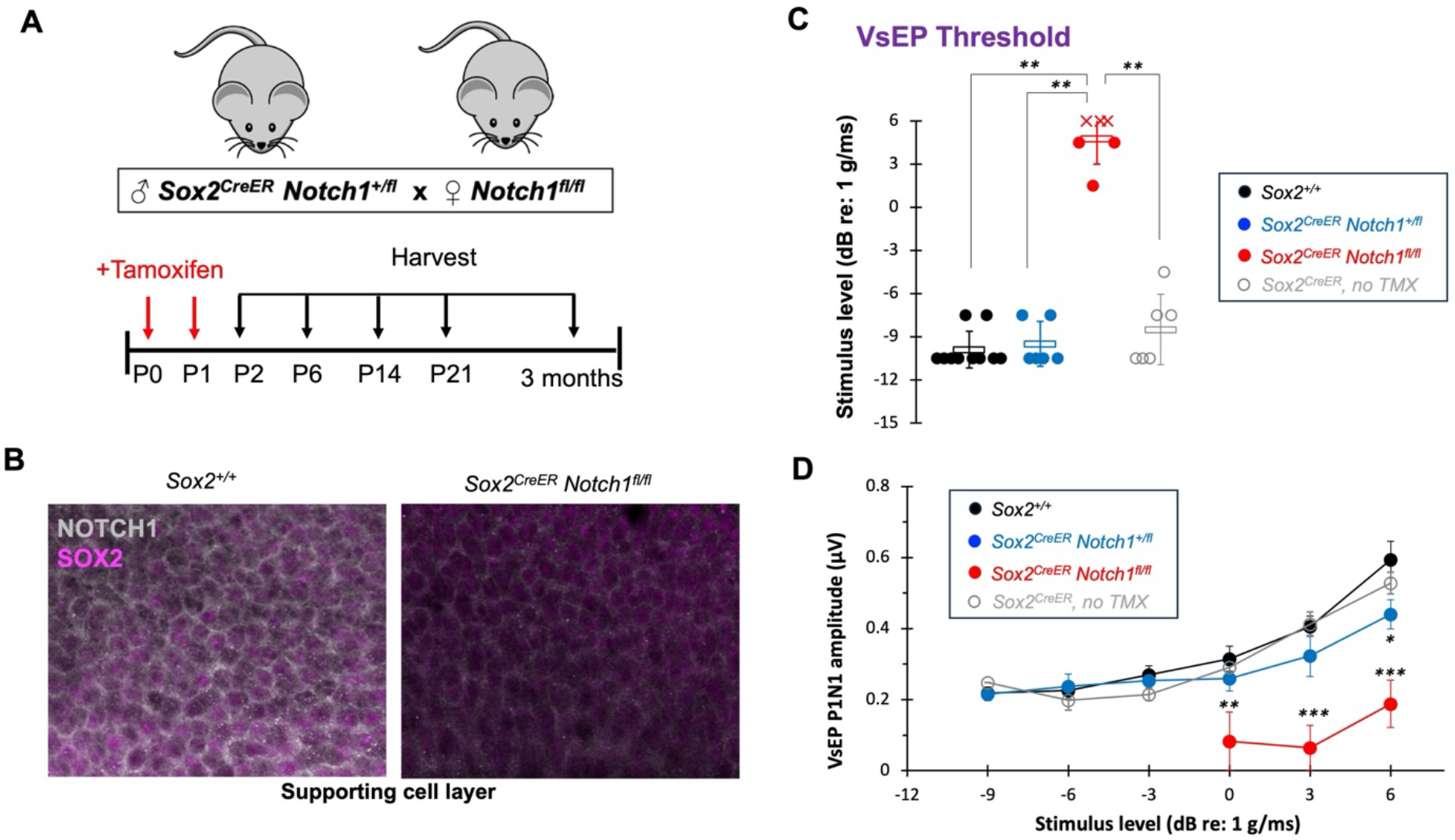
Deletion of *Notch1* at birth causes severe vestibular impairment at 3 months. A) Schematic of conditional *Notch1* deletion in the postnatal vestibular system. *Sox2^CreER^* males were crossed to *Notch1^flox/flox^* females and pups were injected at P0 and 20-24 hours later at P1. Animals were harvested at various time-point of interest, up to 3 months of age. B) Immunofluorescence and confocal imaging shows NOTCH1 is expressed in all vestibular supporting cells, which is greatly reduced in *Sox2^CreER^ Notch1^flox/flox^* utricles by postnatal day 6 (P6). C) Vestibular sensory evoked potential (VsEP) recordings indicate that vestibular function is significantly impaired in Notch1 mutants (red circles, n=6) compared to tamoxifen-injected control (black circle, n=10) and heterozygous (blue circle, n=6) littermates and uninjected Sox2CreER animals (green circles, n=6). “X” symbols denote “no response.” Neither one copy of Notch1 (blue circles) or Sox2 haploinsuffiency (green circles) affects VsEP thresholds. D) *Notch1* mutants have significantly lower P1N1 amplitudes at stimulus levels above 0dB, and at 6dB, heterozygous animals also have a significantly lower amplitude compared to *Sox2^+/+^* controls. Only significant p-values are shown on graphs. *: *p*<0.5; **: *p*<0.01; ***: *p*<0.001.

We first wanted to see if conditional deletion of *Notch1* at birth had any effect on normal vestibular function. To examine this, we performed vestibular sensory evoked potential (VsEP) recordings, which are a direct test of functional output of linear head movements (i.e, the utricle and saccule). We found that compared to both uninjected *CreER* (CreER no TMX), control (*Sox2^+/+^*), and heterozygous (*Sox2^CreER^Notch1^+/fl^*) animals, *Sox2^CreER^Notch1^fl/fl^* mice (*Notch1* mutants) had significantly elevated or undetectable VsEP responses (Figure 1C,D). While *Sox2^CreER^Notch1^+/fl^* heterozygotes did not have a significant increase in VsEP threshold, the average P1-N1 amplitude at the maximum stimulus level of + 6dB was significantly lower than controls (Figure 1D). These data show that deletion of *Notch1* during vestibular maturation leads to compromised macular function.

### Notch1 mutants exhibit behaviors related to vestibular dysfunction

Compared to control (*Sox2^+/+^*) and heterozygous (*Sox2^CreER^Notch1^+/fl^*) littermates, *Sox2^CreER^Notch1^fl/fl^* mice did not exhibit overt behaviors often associated with vestibular impairments, such as circling or noticeable head bobbing (Lee et al., 2002; Somma et al., 2012). To assess whether deletion of *Notch1* affected more subtle vestibular behaviors, we performed an array of other behavioral assessments when the mice were 2-3 months old, including open field, balance beam and swim tests.

Open field tests have been used as an indication of anxiety levels in mice, as well as a way to detect vestibular deficits (Kraeuter et al., 2019). We tracked control, heterozygous and *Notch1* mutant mice for 15 minutes in an open circular arena and compared average velocity and distance traveled between these groups, as well as with uninjected controls (Supplemental Figure 1). We found that overall, *Sox2^CreER^Notch1^fl/fl^* mice did not travel significantly more distance (Supplemental Figure 1D) or have a higher velocity (Supplemental Figure 1E) when compared to all other groups. However, looking at the overall movement of the animals during the open field test, we found that while control mice explored the entirety of the circular arena, *Sox2^CreER^Notch1^fl/fl^* mice seemed to spend more time exploring the central area of the arena (Supplemental Figure 1A-C). They also made significantly more tight circles around their body axis in the central area while exploring the arena than controls (Supplemental Figure 1F), a behavior less-often observed in control mice and suggestive of vestibular dysfunction.

We also assessed dysfunction in the vestibular system using the balance beam test (Luong et al., 2011). Mice were trained for two consecutive days on a 20mm-wide bar, and then challenged with a 9mm-wide bar on the third day. Whereas *Sox2^+/+^* and uninjected controls walked across the 9mm bar without difficulty (Figure 2A,B), *Sox2^CreER^Notch1^fl/fl^* mice took significantly longer to traverse the same 9mm balance beam (Figure 2A,B), and some animals did not complete the task in the allotted two minutes. Additionally, all *Sox2^CreER^Notch1^fl/fl^* mice clearly demonstrated behaviors associated with imbalance that were not seen in controls including wrapping their tails around the bar for stability, slipping of their back legs on the bar, and a head tilt while walking (Figure 2B).

**Figure 2.**
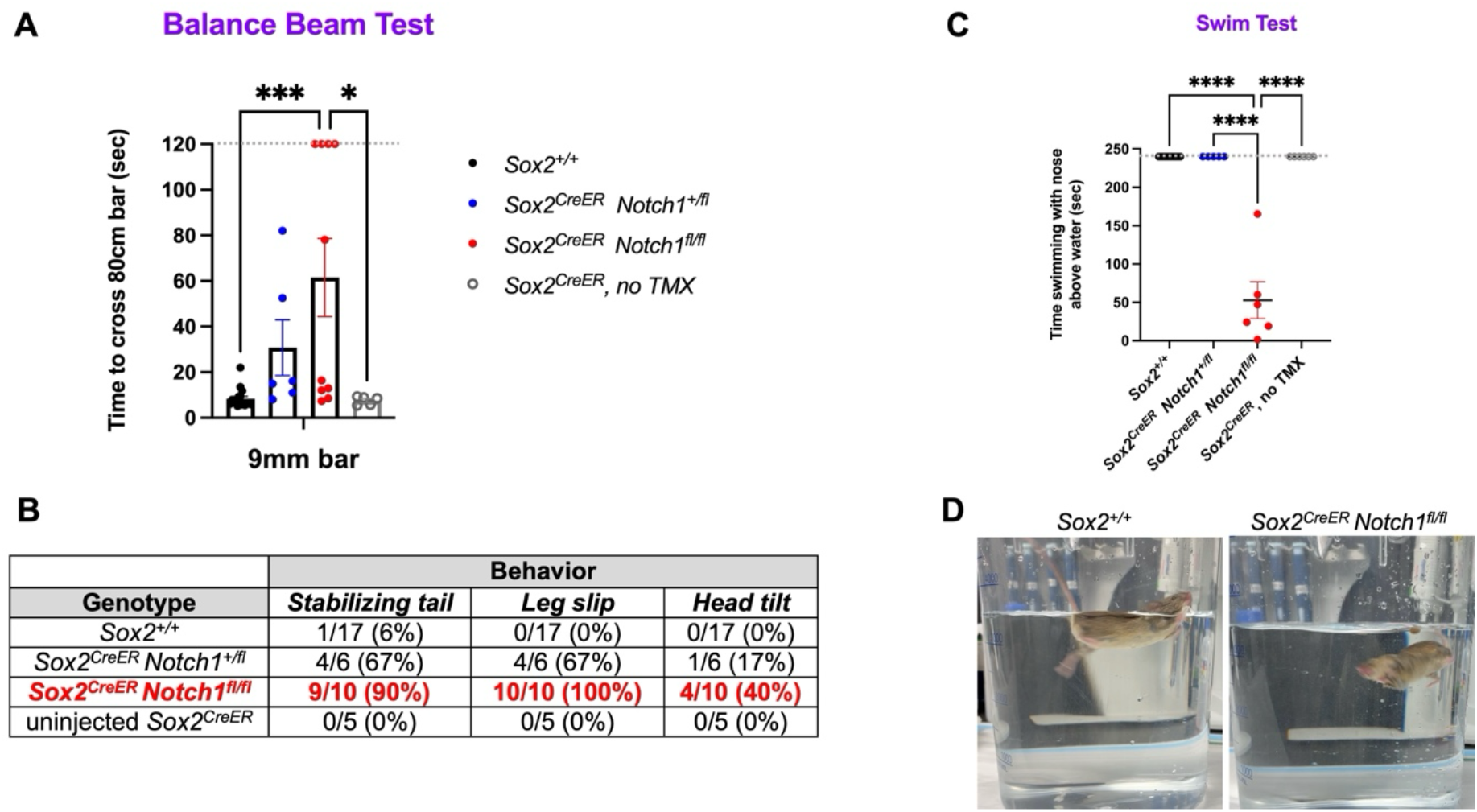
Notch1 mutants exhibit vestibular problems in the balance beam and swimming behavior tests. A) Time (in seconds) it took for animals to cross the 9mm balance bar on the third day of testing. *Sox2^CreER^ Notch1^flox/flox^* animals took significantly longer to cross the bar than both injected and uninjected controls. B) Quantification of behaviors observed during the balance beam test that are associated with balance issues. All *Sox2^CreER^ Notch1^flox/flox^* animals exhibited one or more of these behaviors. C) *Notch1* mutants failed to keep their nose and head above water while swimming. D). Representative frame from swimming video demonstrating a control animal swimming with its head above water (left) and a *Notch1* mutant unable to stay above water (right). Dotted lines are the maximum amount of time allotted in the behavior test. Only significant p-values are shown on graphs. *: *p*<0.5; ***: *p*<0.001, ****: *p*<0.0001.

The final behavioral test we performed was the swim test, where animals were monitored for their ability to keep their head above water for 4 minutes while swimming. We found that while all control and heterozygous mice were able to keep their heads above the water for the duration of the swim test, none of the *Sox2^CreER^Notch1^fl/lf^* mice completed the 4 minute swim test, only being able to keep their heads above the water for an average of 53 seconds before their head went underwater (Figure 2C,D). While trying to swim, *Sox2^CreER^Notch1^fl/fl^* mice exhibited abnormal swimming behavior compared to controls, which included attempting to climb up the container walls, periods of immobility, and bobbing their heads in and out of the water. In severe cases, the animals exhibited body spiraling underwater. Together, these behavior tests support our VsEP results that *Notch1* deletion during sensory maturation results in vestibular dysfunction by 2-3 months of age.

### Notch1 mutants have significantly fewer supporting cells in adult vestibular organs

Next, we investigated what was happening at the cellular level in the vestibular sensory organs that may cause the impaired VsEP responses and behaviors described above. We previously reported that in the cochlea, *Notch1* is required for survival of the sensory outer supporting cells, which include the Deiters cells and outer pillar cells (Heffer et al., 2023); we hypothesized that *Notch1* may function similarly in the maturing vestibular organs. We found that in *Sox2^CreER^Notch1^fl/fl^* mice at 2-3 months, there was also a significant loss of supporting cells throughout the utricle, as seen by a reduction in the number of SOX9+ cells in *Notch1* mutants (Figure 3). Quantification of supporting cell number in the striolar and lateral extrastriolar regions of the utricle showed that there was a significant decrease of SOX9-positive cells in *Notch1* mutants: there was, on average, an 80% reduction in the striola region and 60% reduction in the extrastriolar regions (Figure 3D). Immunohistochemical analysis of vestibular sections in adult mice showed similar supporting cell loss in the saccule and cristae (Supplemental Figure 2). In some regions, there were areas devoid of cells where the supporting cell layer would typically be located (Supplemental Figure 2E, asterisks).

**Figure 3.**
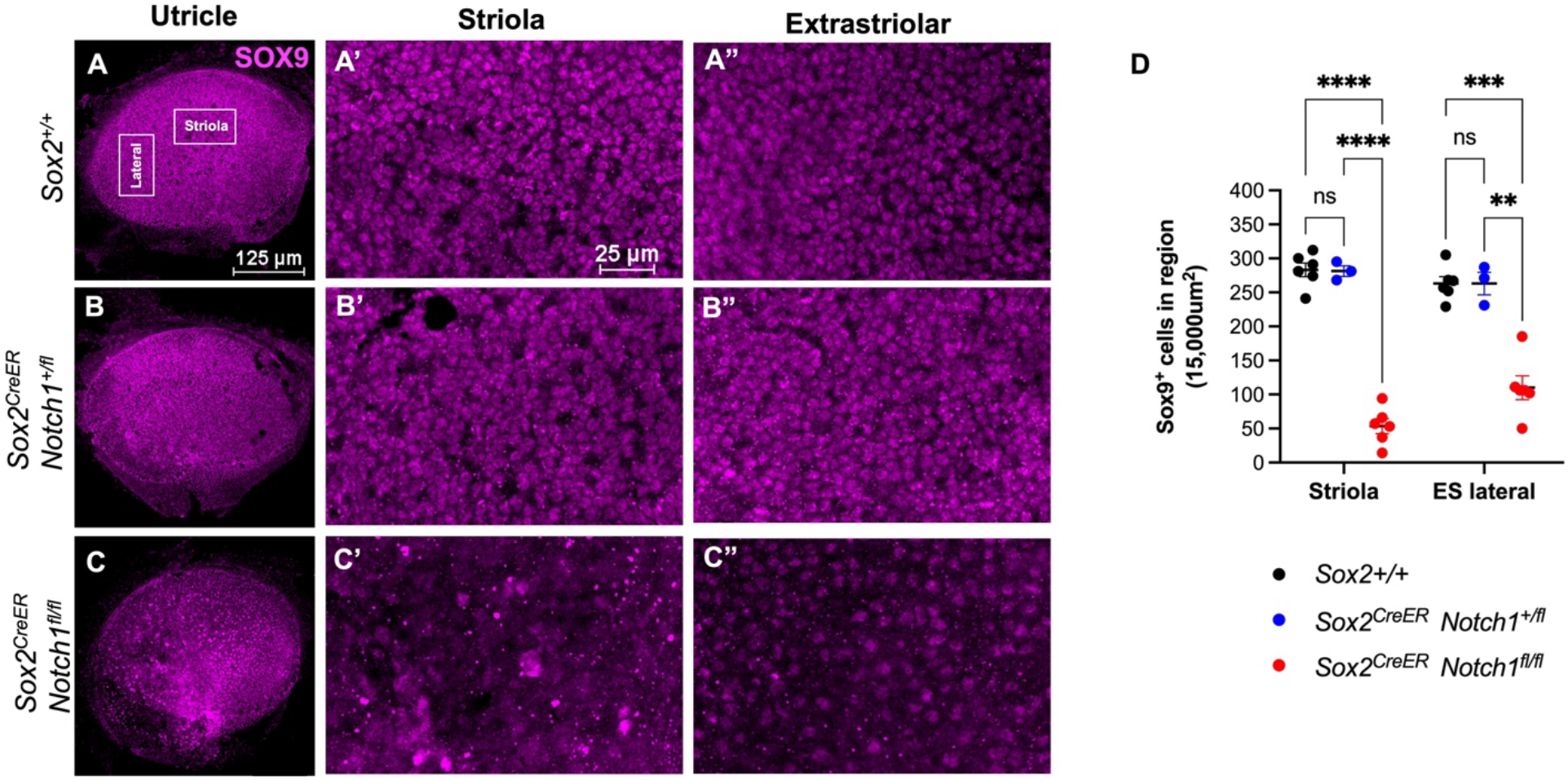
*Sox2^CreER^ Notch1^flox/flox^* animals exhibit supporting cell loss throughout the utricle. A-C) SOX9 staining of utricles of control (A-A”), heterozygous (B-B”) and *Notch1* mutant (C-C”) utricles at 6 weeks. D) Quantification of SOX9+ cells in the striolar and extrastriolar regions show that *Sox2^CreER^ Notch1^flox/flox^* animals have significantly less SOX9+ cells that both *Sox2^+/+^* and *Sox2^CreER^ Notch1^+/flox^* heterozygous animals. **: p<0.01; ***: p<0.001, ****: p<0.0001.

### Vestibular supporting cells are not dying, but are transdifferentiating into type II hair cells

We next examined what was happening to the supporting cells in *Sox2^CreER^Notch1^fl/fl^* mice. We hypothesized that the supporting cells could either be undergoing cell death similar to the cochlea (Heffer et al., 2023), or transdifferentiating into hair cells, suggesting a continued role of Notch in lateral inhibition postnatally. We first immunostained utricles with cleaved-Caspase3 (cCASP3) to look for cells undergoing apoptosis. We found that compared to controls, at no time-point during the first week after tamoxifen treatment did *Sox2^CreER^Notch1^fl/fl^* utricles show any significant difference in the number of cCASP3+ supporting cells (Figure 4A,B) despite decreasing in number, suggesting these cells were not undergoing cell death.

**Figure 4.**
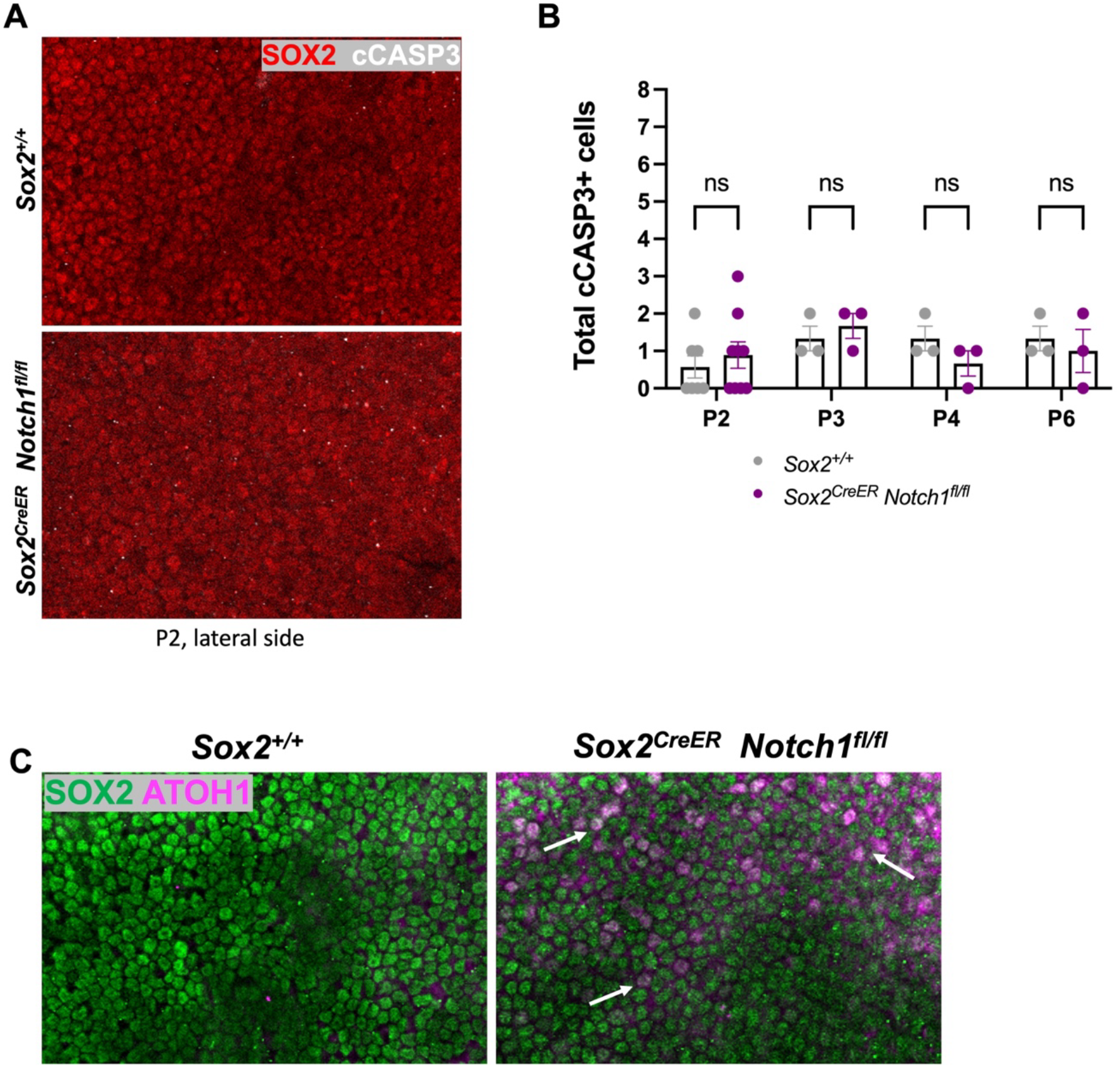
Supporting cells are not dying after *Notch1* deletion, but are converting to hair cells. A,B) cCASP3 staining of supporting cells during the first week after deletion revealed no significant differences in the number of apoptotic cells in *Sox2^+/+^* controls compared to *Sox2^CreER^ Notch1^flox/flox^* mutants. C) Immunostaining with ATOH1 showed no ATOH1+ supporting cells (co-stained with SOX2) in controls at P2, but several ATOH+ supporting cells in *Sox2^CreER^ Notch1^flox/flox^* utricles (white arrows).

We next hypothesized that NOTCH1 may be important during maturation to prevent supporting cells from converting into hair cells, consistent with its role in lateral inhibition. To examine this possibility, we looked for expression of the early hair cell marker ATOH1 in supporting cells. At P2, while there was no ATOH1 expression observed in supporting cells in controls, there were numerous ATOH1+ supporting cells throughout the utricle in *Notch1* mutants (Figure 4C). Together, these results suggest that the supporting cells in the vestibule are not dying after loss of Notch signaling, but rather may be turning into hair cells, supporting a role for NOTCH1 in maintaining supporting cells during vestibular maturation.

In the vestibular epithelia, there are two main types of hair cells – type I and type II – that vary in morphology and gene expression (Burns and Stone, 2017; McInturff et al., 2018). Specifically, type I hair cells are more flask-shaped, with their nuclei located closer to the supporting cells. Type II hair cells are cylindrical in shape and their nuclei are located more apically. We wanted to examine whether these ATOH1+ supporting cells become type I or type II hair cells, and also if they survive to adulthood. We co-stained adult utricles from *Sox2^+/+^* control, *Sox2^CreER^Notch1^+/fl^* heterozygotes and *Sox2^CreER^Notch1^fl/fl^* mutants with SPP1, which labels mature type I hair cells (McInturff et al., 2018), and SOX2, which labels mature type II hair cells (Stone et al., 2021). Quantification of hair cells in the lateral extrastriolar region revealed that while there is no significant difference in the number of type I hair cells in the utricle (Figure 5A-D), *Sox2^CreER^Notch1^fl/fl^* utricles had 2-3 times as many type II hair cells in the same region (Figure 5E-H). Further co-staining of the SOX2+ hair cells in *Notch1* mutants with the mature type II hair cell markers ANXA4 and CALB2 confirmed that the new hair cells are in fact type II hair cells, and also that they are able to mature after arising from supporting cells (Figure 5I-L). These results indicate that ATOH1-positive supporting cells, detected at P2, converted into type II hair cells, and that these hair cells were able to survive to adulthood despite having very few supporting cells (Figure 3). Additionally, examining SOX2 expression in other vestibular end organs (saccule and crista) demonstrated that other vestibular organs also show extra SOX2+ hair cells after *Notch1* deletion, suggesting a similar conversion of supporting cells to type II hair cells in all vestibular organs (Supplemental Figure 2).

**Figure 5.**
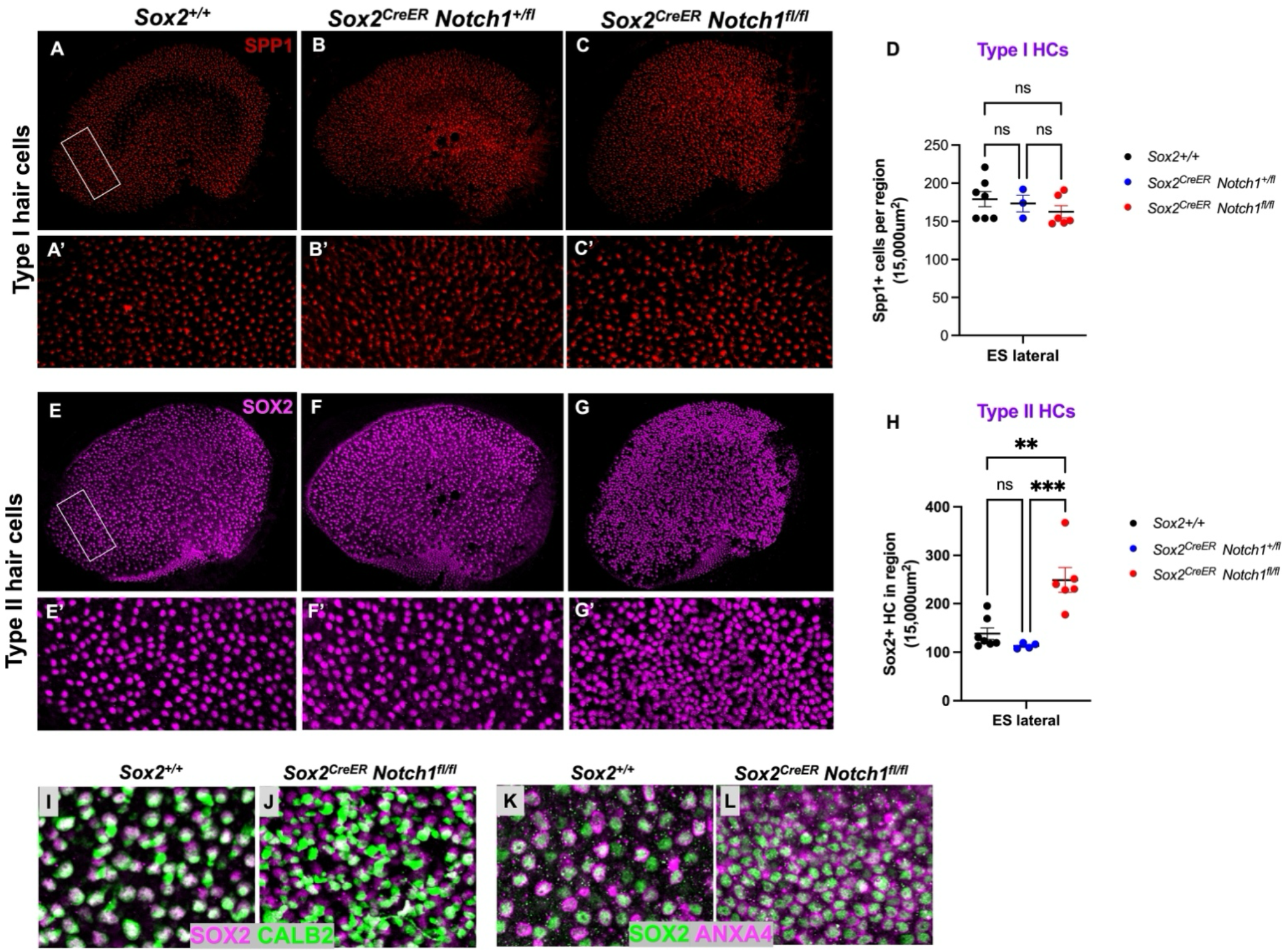
*Notch1* mutants have supernumerary type II hair cells. A-C) SPP1 antibody staining of control (A), heterozygous (B) and *Notch1* mutant (C) adult utricles 6 weeks after *Notch1* deletion. D) The number of SPP1 cells was counted in a 10×15um region of the extrastriola (A’-C’), showing no difference in type I hair cell counts for these genotypes. E-G) SOX2 antibody staining of type II hair cells in the control (E), heterozygous (F) and *Notch1* mutant (G) utricles (same utricles for Type I HC counting). H) SOX2+ hair cells were quantified in the same extrastriolar region as type I hair cells. The SOX2+ nuclei of the type II hair cells of *Notch1* mutants also stained positive for CALB2 (I-J) and ANXA4 (K-L) at P21, showing maturation after conversion. ns: not significant; **: p<0.01; ***: p<0.001.

### Notch1 mutants have a reduced number of specialized type I hair cells in the striola

In addition to the SPP1-expressing type I hair cells found throughout the utricle, there is another population of specialized type I hair cells found only in the striolar region that express the protein Oncomodulin (OCM) throughout the cell body (Hoffman et al., 2018). Since the VsEP signal originates from these specialized type I hair cells in the striola of the maculae sensory organs (Ono et al., 2020; Kim et al., 2022), and the finding that the VsEP responses of our *Sox2^CreER^Notch1^flox^* animals were severely compromised (Figure 1), we hypothesized that there may be abnormalities with the OCM-expressing type I hair cells in the striola of *Notch1* mutants. We immunostained and imaged adult utricles with OCM and found that compared to *Sox2^+/+^* control animals, both heterozygous *Sox2^CreER^Notch1^+/fl^* and mutant *Sox2^CreER^Notch1^fl/fl^* animals had significantly fewer OCM+ cells compared to control utricles (Figure 6A-D), and that these cells appeared to be spread more diffusely across the utricle in the *Notch1* mutants (Figure 6C). Additionally, whereas SPP1-expressing type I hair cells are less numerous in the striola where the OCM-expressing cells are, we find more SPP1-expressing cells around the remaining OCM+ cells in *Notch1* mutant utricular maculae (Supplemental Figure 3).

**Figure 6.**
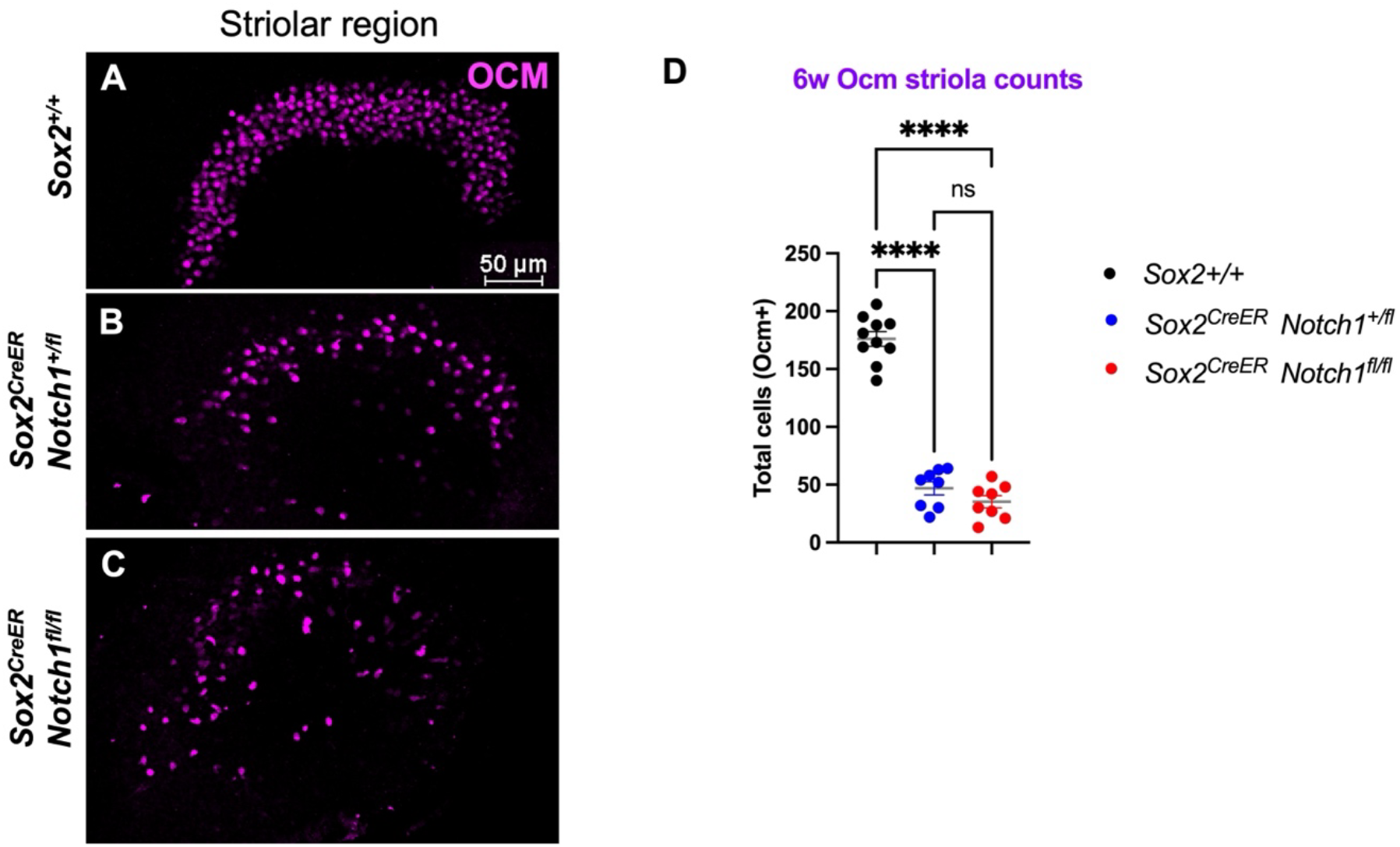
The striola region is abnormal in *Notch1* mutants at 6 weeks. A-C) OCM antibody staining of specialized type I hair cells in the striola in control (A), heterozygous (B) and *Notch1* mutant (C) utricles. D) Quantification of the OCM+ cells in these utricles shows that both heterozygous and *Notch1* mutant utricles have significantly less OCM+ cells at 6 weeks, though these cells are more spread out in the *Sox2^CreER^ Notch1^fl/fl^*animals. ns: not significant; ****: p<0.0001.

### Innervation of hair cells in Sox2^CreER^Notch1^fl/fl^ animals is abnormal

Since *Sox2^CreER^Notch1^+/fl^* and *Sox2^CreER^Notch1^fl/fl^* animals had a similar number of OCM+ cells in the utricle at 6 weeks but *Sox2^CreER^Notch1^fl/fl^* animals had largely absent VsEP responses, these results suggest that there may be a problem with the afferent innervation of striolar hair cells in *Notch1* mutant animals. Normally, groups of two or three type I (OCM+) hair cells in the striola are innervated by a single afferent with complex calyceal endings, which can be visualized with Calbindin2 (CALB2) antibody staining. To examine this, we immunostained utricles with CALB2 and the neuronal marker TUJ, which co-stains complex calyces as well as afferent boutons of type II hair cells and simple calyces that surround type I hair cells. We found that in control utricles, OCM+ cells are surrounded by CALB2+ and TUJ+ calyces in groups of two and three throughout the striolar region (Figure 7J,M,P; arrows, arrowheads). In heterozygous utricles, while there are fewer OCM+ cells, the cells that remain still form complex calyces, mostly in groups of 2, which are surrounded by CALB2+/TUJ+ innervation (Figure 7K,N,Q; arrowheads). In contrast *Notch1-*deficient utricles displayed very few CALB2-positive calyces, most of which do not form complex calyces and have weaker CALB2 staining (Figure 7L,O,R; asterisks). This can be observed clearly in images that show both CALB2 (green) and TUJ (red) staining (Fig 7P-Q), in which complex calyces show dual labeling of CALB2 and TUJ1 (yellow), whereas very few co-labeled structures are found in *Notch1*-deficient utricles (Fig 7R) Interestingly, TUJ staining is still found around all OCM+ cells (Figure 7O), indicating they are still innervated, but the innervation resembles that of non-OCM+ type I cells.

**Figure 7.**
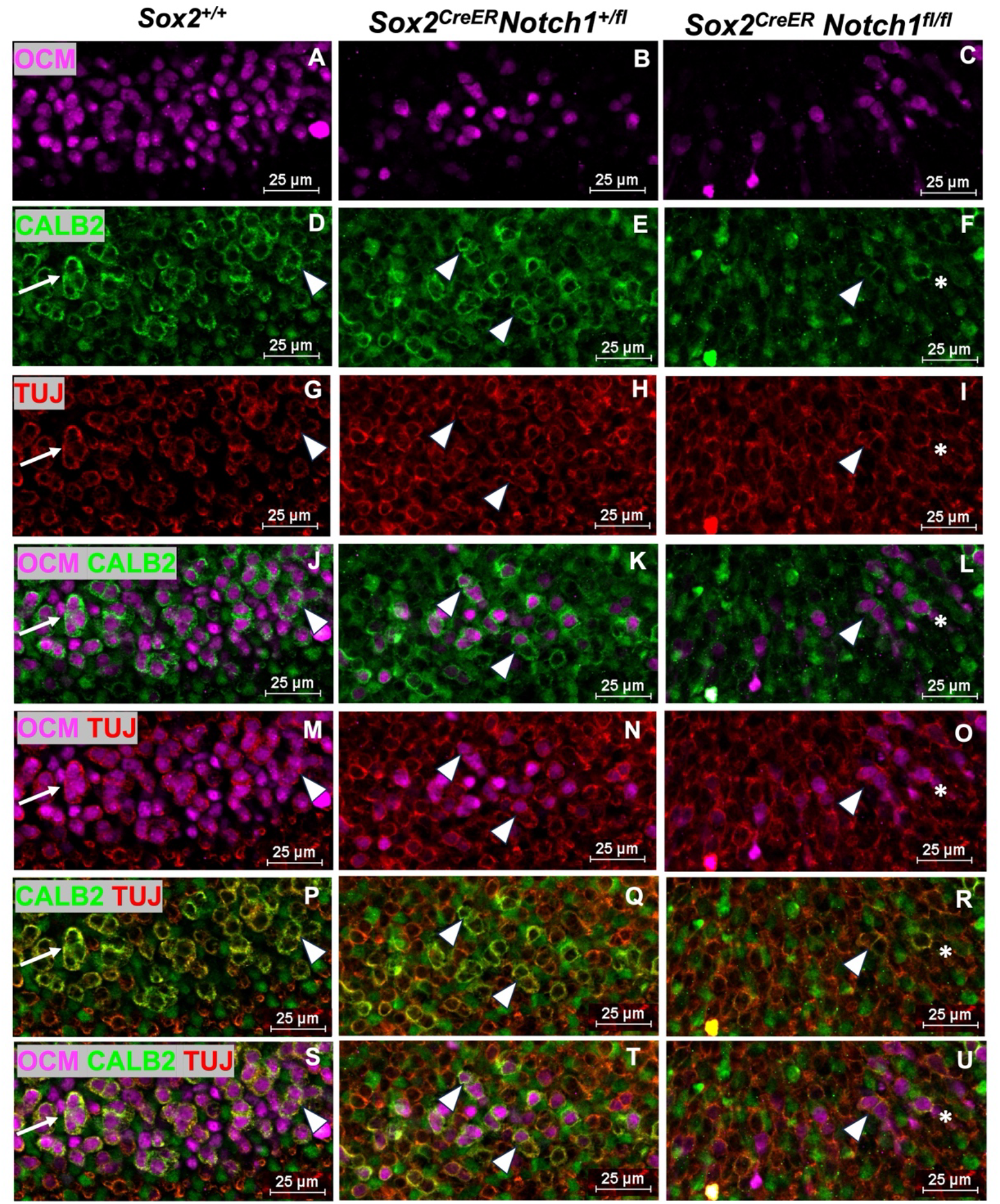
Innervation of the striola in is abnormal in *Notch1* mutants at 6 weeks. Left column (A,C,G,J,M,P,S): In control *Sox^+/+^* animals, OCM+ cells in the striola (A) are mostly innervated as complex calyces that form groups of 2 (arrowheads) or 3 (arrows) and stain positive for CALB2 and TUJ (D,G,P,S). Middle column (B,E,H,K,N,Q,T): heterozygous animals have fewer OCM cells in the striola (B), and these cells are mostly in groups of 2 (E,H,Q,T). Right column (C,F,I,L,O,R,U): Cells that stain positive for OCM in *Notch1* mutants are more spread out and are occasionally found in groups of 2 (arrowhead), but normally as single cells (asterisk). CALB2 staining is weaker (F,L) the OCM cells that remain are more often associated with TUJ rather than CALB2 (I,L,O).

While expression of OCM in striolar type I hair cells can be detected as early as E18.5 and immature neurons and calyces can be detected as early as postnatal day 0, the more complex calyces aren’t mature until the second postnatal week (Burns and Stone, 2017). We sought to determine whether the OCM+ cells in *Notch1* mutants innervated properly and developed complex calyces during maturation by co-staining P14 utricles with OCM, CALB2 and TUJ (Supplemental Figure 4). In *Sox2^+/+^* controls, many OCM+ cells stained positive for both CALB2 and TUJ at P14 (Supplemental Figure 4A). *Sox2^CreER^Notch1^fl/fl^* utricles did not have this same co-localization of OCM, CALB2 and TUJ, but rather the CALB2 staining appeared as punctate dots and did not overlap with TUJ (Supplemental Figure 4B); much of the CALB2 staining we observe is likely due to the type II hair cell staining in this area. We examined utricular cross-sections at this same time and found similar results: while many OCM+ cells were surrounded with calyces staining positive for CALB2 and TUJ, there was no co-localization of CALB2 and TUJ around the specialized type I hair cells that were present (Supplemental Figure 4C,D). In fact, CALB2 protein seemed to form aggregates within the hair cell layer between type II hair cells (Supplemental Figure 4D’’). Similar to what we saw at 6 weeks, some OCM+ cells had TUJ+ calyces but not CALB2, which could be still be immature or alternatively, more reminiscent of the non-specialized type I hair cells (Supplemental Figure 4D). Additionally, whereas control OCM expression was found throughout the hair cell body, OCM expression in *Notch1* mutants appears to localized more to the apical tip of the hair cell and not be strongly expressed in the hair cell body (Supplemental Figure 4D’).

### Innervation of hair cells in the extrastiolar region is also abnormal

Since innervation of hair cells in the striola was abnormal in *Sox2^CreER^Notch1^fl/fl^* utricles, we looked to see if the type I and type II hair cells in the extrastriolar region were innervated properly, and whether the new type II hair cells that arose after deletion of *Notch1* became innervated. Antibody staining of neuron terminals with TUJ showed that there was a similar distribution of TUJ+ neurons present in control and heterozygous utricles, but in *Notch1* mutant utricles the staining pattern was very disorganized and had partial calyces and boutons with more punctate or incomplete regions (Figure 8A-C). Further examination of hair cell innervation through utricle cross-section analysis revealed that in controls, type I hair cells were surrounded by a calyx and the nuclei were closer to supporting cells, where type II hair cells were located closer to the apical surface. In *Notch1* mutants, type I hair cells still appeared to have calyces surrounded them, and most type II hair cells also appeared localized to TUJ staining, often having TUJ “branching” points below (Figure 8D’-F’; arrow). However, due to the hair cell layer being less uniform in hair cell distribution in *Notch1* mutants, we cannot be certain if the new hair cells that arise from supporting cells are actually innervated properly.

**Figure 8.**
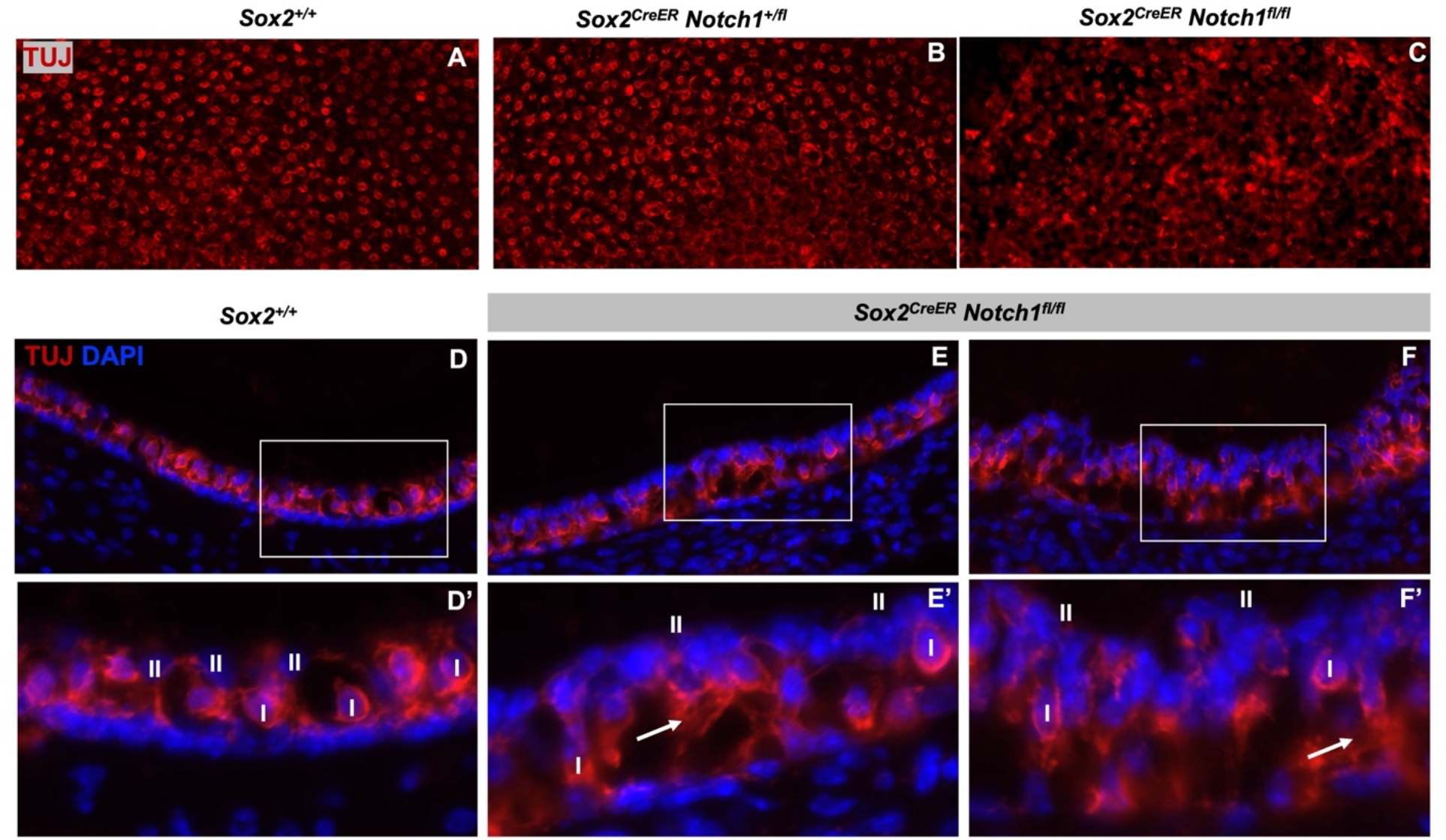
Innervation of extrastriolar hair cells appears abnormal in *Notch1* mutants. Staining utricles by whole mount immunofluorescence with TUJ shows organized innervation of type I and type II hair cells in the extrastriolar region of control (A) and heterozygous (B) adults. In *Notch1* mutants (C) TUJ staining shows disorganized TUJ staining in the extrastriolar region, with areas of aggregation (brighter areas). D-F) Cross sections of 6 week utricles conform this dis-organization of innervation. In controls (D), the type I hair cells surrounded by simple calyces and the nuclei are located closer to the supporting cells (row of DAPI cells) and type II hair cell nuclei are located closer to the apical side of the sensory epithelium (D,D’). E,F) The innervation of *Notch1* mutants is much more disorganized, with the positions of the type I and type II hair cells less defined (E’,F’). Additionally, *Notch1* mutants have regions of higher density of TUJ staining (arrows), which are likely points where new neurons branch off to innervate the new hair cells.

## Discussion

Here, we explored the role of Notch signaling during maturation of the vestibular system by conditionally deleting *Notch1* at birth in the postnatal maturing supporting cells of the balance organs. Histological analysis showed a dramatic loss of supporting cells; however, unlike in the auditory system (Heffer et al., 2023), supporting cells did not undergo cell death but instead transdifferentiated into type II hair cells. Surprisingly, we found that adult NOTCH1-deficient mice exhibited no obvious behaviors normally associated with vestibular defects, including circling and head bobbing. However, these mutant mice demonstrated difficulties traversing a balance beam, difficulties with swimming, and differences in open field activity indicating there was some vestibular impairment. In addition, VsEPs, a test of macular function using linear head motions, was absent in *Notch1*-deleted animals. Interestingly, *Notch1* mutants showed a loss of complex calyces that normally are present surrounding the type I OCM+ hair cells, correlating to loss of striolar function. Together, these results indicate that vestibular supporting cells are more plastic than auditory supporting cells and retain the ability to transdifferentiate into hair cells. Surprisingly, despite the lack of supporting cells and excess hair cells, the majority of the hair cells survive into adulthood and show innervation.

### Hair cell survival despite reduction of supporting cells

Previously we have shown that in the cochlea, loss of NOTCH1 causes rapid death of the supporting cells within 24-48 hours. However, outer hair cells and some inner hair cells are not lost until about two weeks later, around P14 (Heffer et al., 2023). This is consistent with studies in which the supporting cells were deleted using diphtheria toxin (DTA), leading to hair cell loss several weeks later (Mellado Lagarde et al., 2013). These results indicate that in the cochlea, survival of the supporting cells is critical for the survival of hair cells. In contrast, in mammalian vestibular organs, supporting cells (at the least the full contingent) do not appear to be critical for hair cell survival, as we observe hair cells across the vestibular sensory organs in adults (Figure 5). Hair cell death following loss of supporting cells has also been documented in non-mammalian species. For example, in zebrafish, mutation of the Notch-related gene *mind bomb* leads to a dramatic conversion of supporting cells to hair cells during development, resulting in rapid loss of the hair cells, presumably due to lack of supporting cells (Haddon et al., 1999). In birds, after hair cell ablation, supporting cells convert directly into new hair cells and are ejected from the epithelia within a few days after conversion (Mangiardi et al., 2004). In the mammalian vestibular system, our data suggest that 20-40% of the normal complement of supporting cells is sufficient for maintaining hair cell survival, which, may be important for future therapies that target supporting cells for hair cell replacement.

### Cellular changes in extrastriolar and striolar regions after Notch1 deletion

We show that by six weeks, while there is no increase in type I hair cells in the extrastriolar region, there is an almost 2-fold increase in the number of type II extrastriolar hair cells (Figure 5). These results indicate that the supporting cells are directly transdifferentiating into type II hair cells in the absence of NOTCH1. These results are consistent with experiments examining regeneration in mature utricles after hair cell ablation (Lin et al., 2011; Golub et al., 2012; Bucks et al., 2017). However, the cellular changes in the striolar region are more complex after *Notch1* deletion. Firstly, the loss of SOX9+ supporting cells is more dramatic in the striolar regions as compared to the extrastriolar regions. Specifically, there is about an 80% loss of supporting cells in the striolar regions as compared to about a 60% loss in the extrastriolar regions. Since there is no detectable increase in cleaved-Caspase-3 activity during the time supporting cell loss is observed, we can assume that this loss cannot be attributed to cell death. Interestingly, we also observe about a 60% loss of OCM+ cells, the specialized type I cells in the striola (Figure 6). The OCM+ cells are also not as tightly organized in the NOTCH1-deficient utricles (Supplemental Figure 3), likely due to converting supporting cells. There also appears to be more non-specialized type I cells that are SPP1+ surrounding the OCM+ cells in *Notch1* mutants (Supplemental Figure 3). Thus, it is possible that in the striolar regions, the supporting cells are converting into both type II and non-specialized type I cells (SPP1+). Another possibility is that the OCM+ cells may be converting into SPP1+ type I cells, or possibly into type II cells. In support of this hypothesis, Ono and colleagues (Ono et al., 2020) also observed that a decrease in OCM+ cells in the striolar region was accompanied by an increase in SPP1+ type I hair cells, similar to what we observe in our *Notch1* mutant mice (Supplemental Figure 3). Interestingly, it has been shown that neonatal supporting cells in the striolar region can give rise to both type I hair cells (SPP1+) and type II hair cells (Wang et al., 2015; Wang et al., 2019), indicating there may be specialized supporting cells in this region. Indeed, these studies have shown that some supporting cells in the striolar region upregulate Lgr5, a Wnt-responsive gene, in response to hair cell loss (Wang et al., 2015).

### Innervation defects and the VsEP response

Multiple lines of evidence indicate that VsEPs are generated by calyx-bearing afferents in the striolar region (Jones et al., 2015; Curthoys et al., 2017; Lee et al., 2017; Ono et al., 2020). The specialized type I hair cells (OCM+) in the striolar region are grouped in bundles of two or three and innervated by complex calyces. In the *Notch1*-deficient animals, we detected many fewer complex calyces (Figure 7), even in the remaining OCM+ hair cells. Since we also failed to detect a VsEP response in these animals, this result is consistent with the idea that these OCM+ type I hair cells and their complex calyces are critical for the VsEP response. Interestingly, although we also detected reduced numbers of OCM+ cells in heterozygotes, these animals still showed a VsEP response, though the P1N1 amplitude was reduced at the highest stimulus level of threshold finding protocol (i.e., +6dB re: 1.0 g/ms). We also detected complex calyces using CALB2 staining in the heterozygote utricles. These results indicate that reduced numbers of type I OCM+ hair cells are still able to generate VsEP responses, as long as they are innervated properly.

The loss of complex calyces also indicates that loss of the supporting cells in the NOTCH1-deficient animals likely contributes to defects in innervation in the striolar region. Interestingly, Ono and colleagues found that disruptions in retinoic acid signaling during embryogenesis resulted in a loss of the striola at birth, which correlated with absence of VsEP responses in adults (Ono et al., 2020). Specifically, they reported that the expression of *Cyp26b1*, an enzyme required for retinoic acid degradation, was expressed in the supporting cells of the striola, whereas the enzymes that produce retinoic acid, including *Aldh1a3*, are expressed in both hair cells and supporting cells in the extrastriola regions (Ono et al., 2020). Thus, a reduction of supporting cells would likely phenocopy loss of *Cyp26b1*, leading to higher retinoic acid concentrations throughout the utricle. Therefore, it is likely that loss of the supporting cells in the NOTCH1-deficient animals also disrupts retinoic degradation, and thus the striolar region. However, in contrast to Ono and colleagues, we are disrupting supporting cells after birth, suggesting that supporting cells are required for the maintaince of the striolar region in the neonatal period. Thus, our results indicate that supporting cells are critical for striolar maintainence during sensory maturation in the vestibular regions, likely through maintenance of the retinoic acid gradient.

### Physiological and behavioral defects

While there has been a fair amount of documentation of cellular changes after vestibular damage, few studies have correlated cellular changes with physiological and behavioral changes. Thus, it has been unclear what type of defects lead to decreased vestibular responses and behavioral deficits. In this study, we combined physiological (VsEP) and behavioral (open field, balance beam, and swimming) studies with immunohistochemical studies to try and correlate specific vestibular behaviors with underlying cellular phenotypes. At present, there are very few studies that have looked at what types of behaviors are governed by the striolar region. Other mutants that lack a striolar region do not show typical vestibular deficits such as circling or head bobbing, but do show deficits in the balance beam (Ono et al., 2020). Our NOTCH1-deficient animals, which lack a striolar region also do not show overt vestibular behaviors but show difficulties on the balance beam (Figure 2). However, in contrast to other striolar mutants, *Notch1* mutant animals also struggle to swim, suggesting there are defects outside of the striolar region that govern swimming behavior. Consistent with this observation, we observe abnormal innervation patterns in the extrastriolar regions compared to control and heterozygote animals (Figure 8). Other mutants that have also been reported to have issues swimming properly display problems with hair cell organization outside of the striola, including Emx2 mutants (Ji et al., 2022).

## Conclusions

Our studies indicate that NOTCH1 is required to maintain supporting cell identity during postnatal sensory maturation in the vestibule. Loss of NOTCH1 leads to supporting cell conversion to type II hair cells throughout the vestibular sensory areas. Our data indicate that many fewer supporting cells can still support hair cell survival, which may be important for vestibular restoration approaches that convert supporting cells to hair cells. However, our data show that supporting cells are critical for correct innervation patterns and thus hair cell function in the striolar and extrastriolar regions of utricle, resulting in loss of VsEP responses and balance and swimming problems in their absence.

## Acknowledgements

This work was supported by National Institutes of Health–National Institute on Deafness and Other Communication Disorders Grants R01 DC009250 and R01 DC017767 to A.E.K., R01 DC016974 to J.C.H., and a departmental grant from Research to Prevent Blindness

---

**Supplemental Figure 1.**
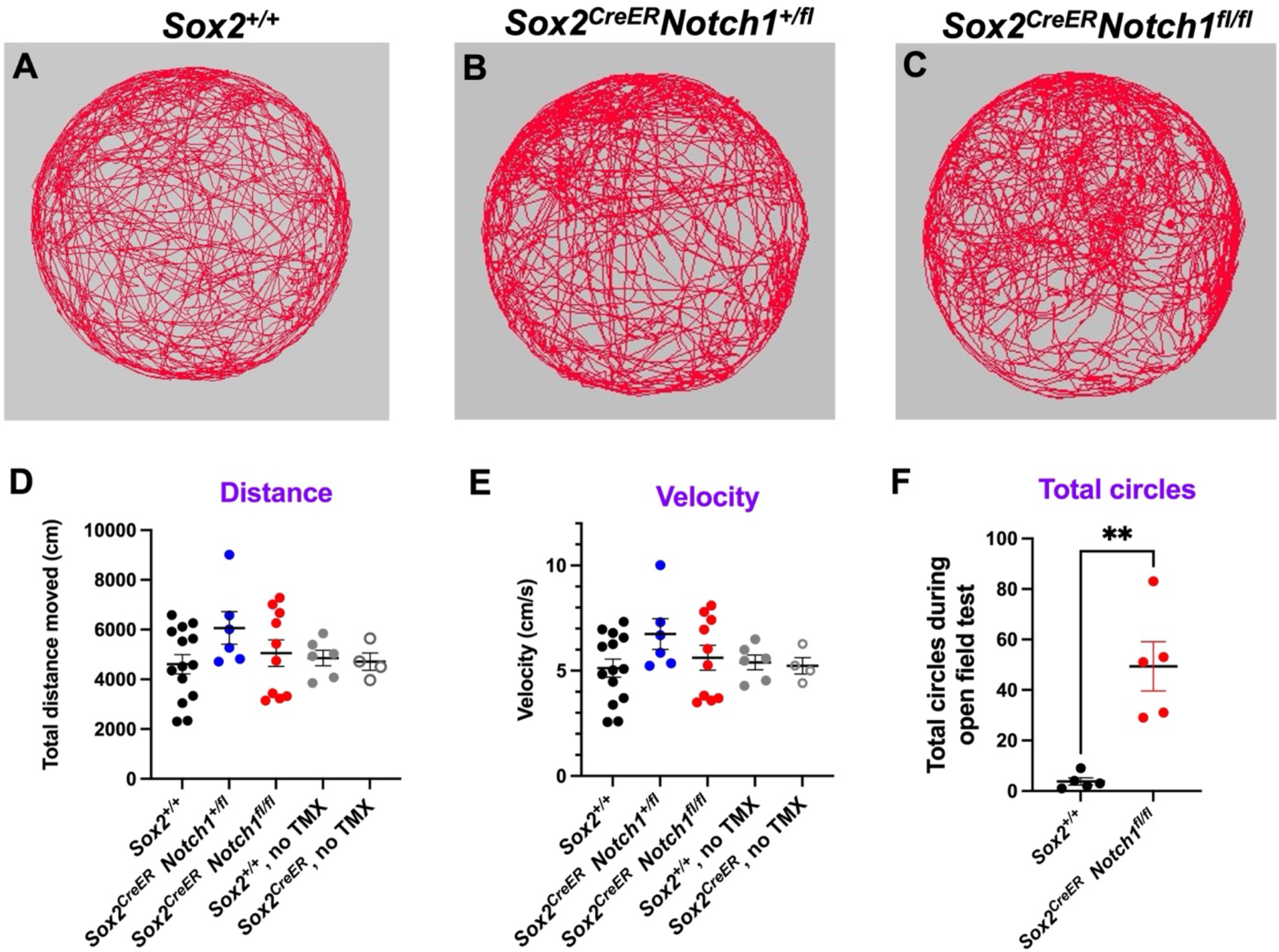
N*o*tch1 mutants display some vestibular dysfunction in open field tests. A-C) Representative trace plots of control (A), heterozygous (B) and *Notch1* mutant (C) mice over the duration of the 15 minute open field test. No groups of mice tested were significantly different in the total distance they traveled during the test (D) or the average velocity they moved at during the test (E). F) *Notch1* mutant mice did spend more time in the center of the arena (C) and also made significantly more tight circles around their body axis in the open field arena, suggestive of vestibular problems (F). Only significant p-values are shown. **: p<0.01

**Supplemental Figure 2.**
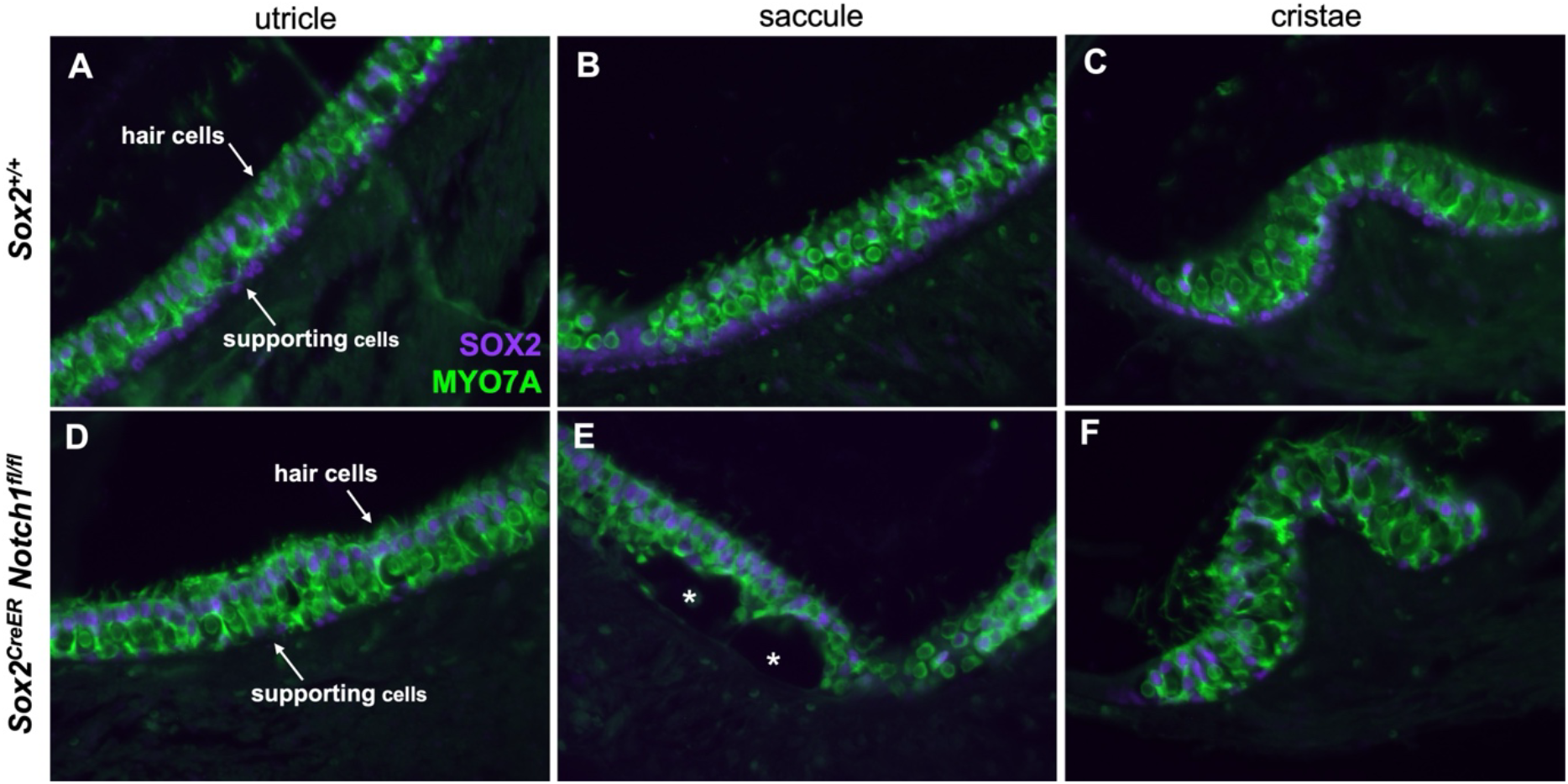
Supporting cell and hair cell phenotypes observed after *Notch1* deletion at P0/P1 in the utricle are also seen in the other vestibular sensory regions at 6 weeks. A-C) Immunostaining of vestibular cross sections show that in control animals, supporting cells stained with a SOX2 antibody (purple) are arranged in a single layer below hair cells stained a MYO7A antibody (green). SOX2 also stains type II hair cells (SOX2+, MYO7A+). D-F). In *Sox2^CreER^ Notch1^flox/flox^*vestibular sensory regions, the majority of supporting cells are missing, and there are more SOX2+ (type II) hair cells. In the saccule (D) of *Notch1* mutants, there were areas where the hair cells were detached from the epithelia, forming “pockets” (white asterisks in E).

**Supplemental Figure 3.**
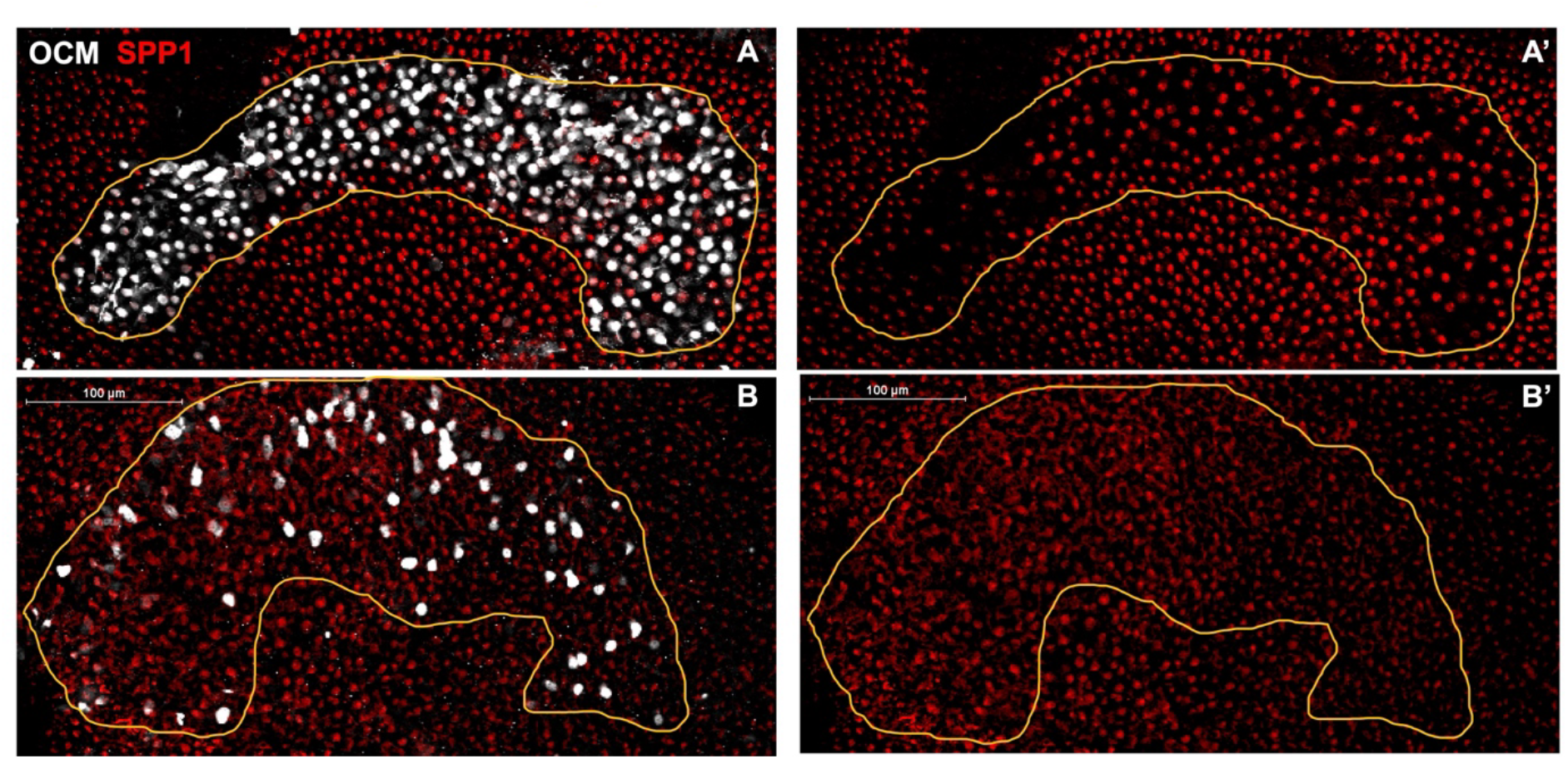
N*o*tch1 mutant utricles have more SPP1+ cells in the presumptive striola at 6 weeks. A,A’) The striolar region of *Sox2^+/+^*animals have a high concentration of OCM-expressing cells, with fewer SPP1+ compared to the surrounding extrastriolar areas. B,B’) *Sox2^CreER^Notch1^fl/fl^* utricles have a more diffuse OCM expression pattern, with SPP1+ type I hair cells found throughout at a similar density to those in surrounding areas. The striolar region was drawn by hand around all OCM+ cells in controls, and a presumptive striolar region was drawn around OCM cells in mutants.

**Supplemental Figure 4.**
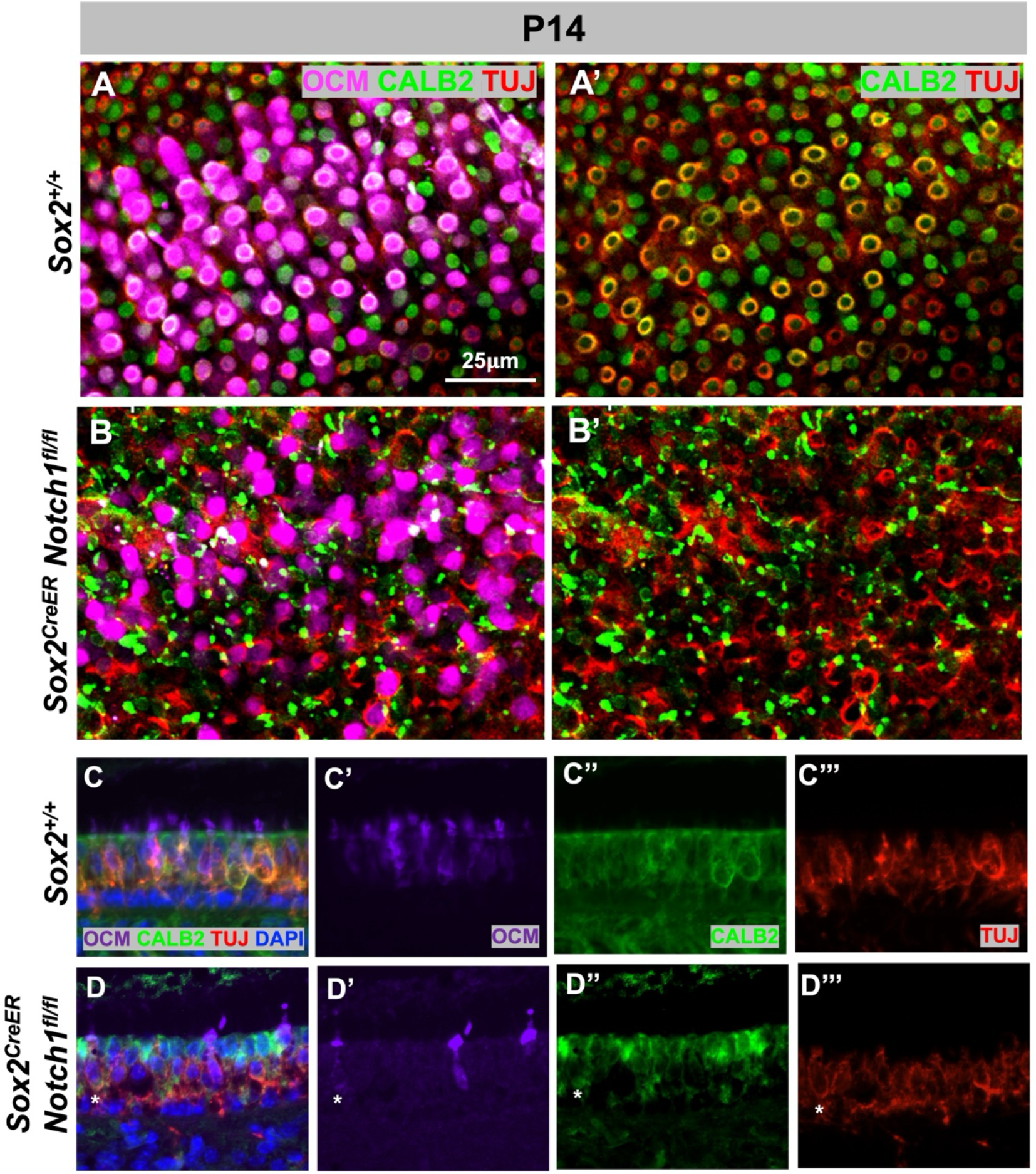
During maturation, innervation of the striola fails to complete in *Notch1* mutants. A,A’) By P14, control utricles show innervation of the complex calyces of OCM cells by co-localization (yellow) of CALB2 (green) and TUJ (red). B,B’) The OCM cells that are present do not show co-localization of CALB2 and TUJ, but rather CALB2 staining is more punctate. C,D) Cross-sections of *Sox2^+/+^* controls (C-C’’’) show that OCM is expressed throughout the cell body (C’) and there are complex calyces that have begun forming (C’’). In *Sox2^CreER^Notch1^fl/fl^*mutants (D-D’’’), OCM is more localized to the hair cell tip (D’) and CALB2 is more aggregated between the hair cells than around OCM cells (D’’). OCM cells are present that show TUJ innervation, but no CALB2 (asterisk), suggesting they are more like type I hair cells found outside the striola (D’’’).

